# Chromosomal capture of beneficial genes drives plasmids towards ecological redundancy

**DOI:** 10.1101/2025.01.21.634075

**Authors:** R. Craig MacLean, Cédric Lood, Rachel M. Wheatley

## Abstract

Plasmids are a ubiquitous feature of bacterial genomes, but the evolutionary forces driving genes to become associated with plasmids are poorly understood. To address this problem, we compared the fitness effects of chromosomal and plasmid genes in the plant symbiont *Rhizobium leguminosarum*. Here we show that plasmids are depleted in beneficial genes compared to the chromosome, and this effect is stronger for ancient plasmids compared to recently acquired plasmids. These findings support the hypothesis that evolution drives beneficial genes to become localized to the bacterial chromosome, resulting in a gradual decay in the ecological value of plasmids. These findings question the ecological importance of plasmids and highlight the challenge of understanding how plasmids persist over the long term.

## Main text

Bacterial genomes are made up of chromosomes and plasmids that replicate independently of the chromosome. Genes are continuously transferred between plasmids and chromosomes, and uncovering the processes that drive genes and phenotypes to be associated with plasmids as opposed to bacterial chromosomes is a fundamental challenge in microbial ecology and evolution(*1-8*).

The dominant view in microbiology is that plasmids play a key role in bacterial adaptation through the horizontal transfer of genes that are beneficial in defined ecological niches(*9-13*), such as genes associated with antibiotic resistance, pathogen virulence, or novel metabolic pathways(*1, 8, 14-17*). However, classic evolutionary models that allow genes to move between plasmids and the chromosome predict that beneficial genes should become associated with chromosomes, as opposed to plasmids, questioning the role of plasmids in bacterial adaptation (*2, 18*). It has been challenging to reconcile these two views of plasmids (*6, 8, 11, 12, 19, 20)* because the relative ecological and evolutionary importance of plasmid genes remains poorly understood beyond the paradigmatic examples highlighted above.

Here we address this problem by systematically measuring the impact of plasmid and chromosomal genes on bacterial fitness in the plant symbiont *Rhizobium leguminosarum*. Wheatley *et al*(*21)* used transposon insertions (*22*) to systematically mutagenize the genome of a strain of *R. leguminosarum* carrying a chromosome and 6 plasmids(*17*). Populations of pooled insertion mutants were then assayed by deep sequencing under conditions that recapitulate the ecology of *Rhizobium* (*23*), including growth in the rhizosphere, root colonisation, nodulation, and bacteroid formation (Figure 1 A,B). The use of fitness assays under natural conditions is a key feature of this data set, given that plasmids are predicted to carry ecologically relevant genes whose effects may be missed in standard lab-culture based measures of bacterial fitness. This experiment uncovered 603 unique genes that were beneficial in either a single niche (specialist genes) or across multiple niches (generalist genes) (Figure 1 B,C).

**Figure 1:**
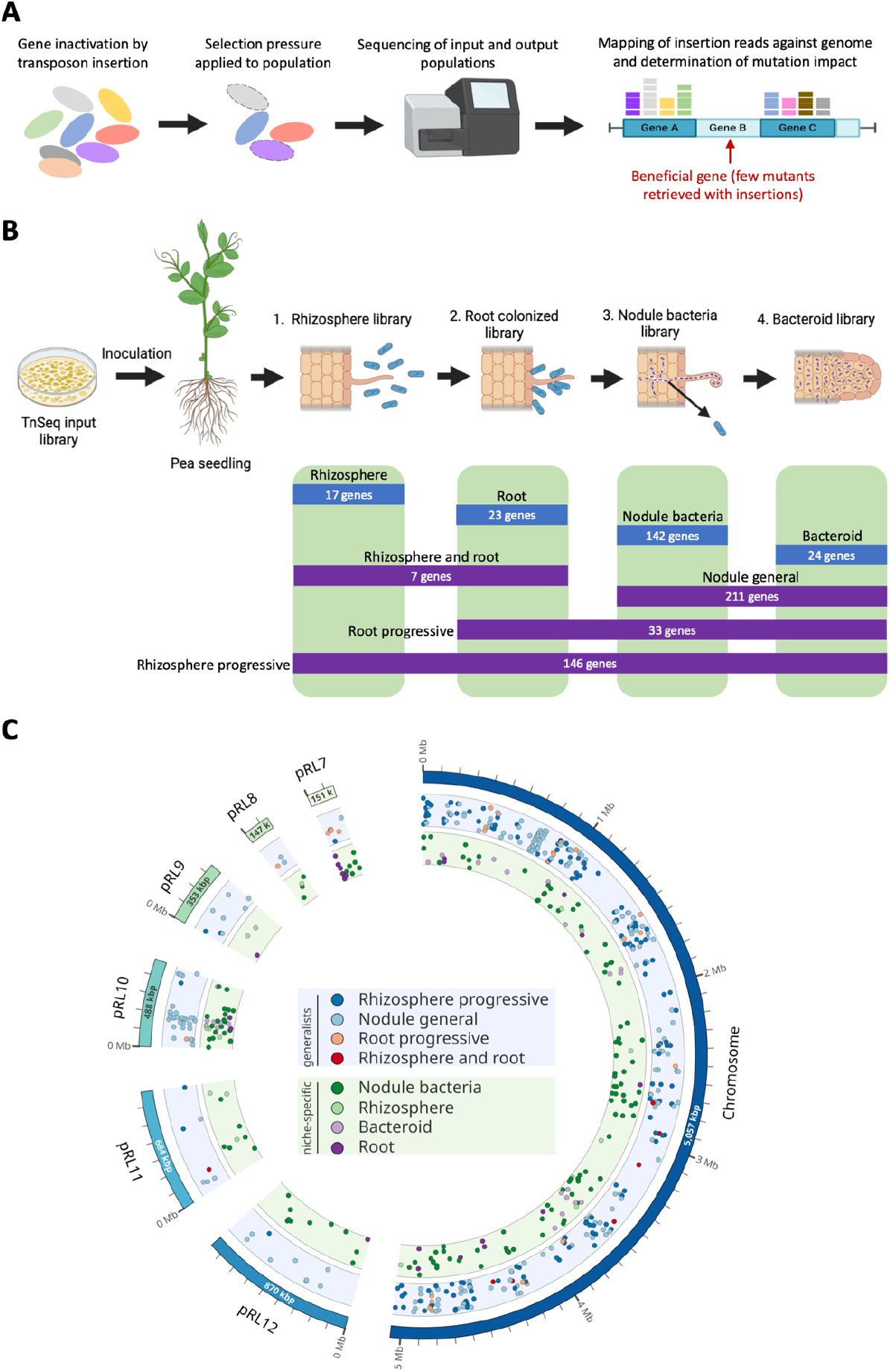
Identifying beneficial genes by Tn-Seq. (**A**) Schematic of a transposon insertion sequencing experiment. First, a mutant library is constructed using transposon insertion to inactivate genes on a genome-wide scale. When a selection pressure is applied to the population, mutants change in frequency in the population depending on the contribution of their mutated gene to fitness under that condition. The resulting populations, including the input library, are then sequenced and mapped back against the genome to determine the position of and frequency of transposon insertion mutants across the genome. Mutants of genes which are important for fitness will fall out of frequency of the population, allowing their identification as beneficial genes. (**B**) Summary of the transposon insertion sequencing experiment previously conducted by Wheatley et al. (*21*) in which a *R. leguminosarum* mutant library was assayed across multiple stages of symbiosis: growth in the rhizosphere, root colonisation, nodule formation, and bacteroid formation. The table indicates the number of genes which were identified to be beneficial across the corresponding niches. Blue blocks indicate the number of genes beneficial in single niches, and purple blocks indicate the number of genes beneficial in multiple niches. This figure was made in biorender. (**C**) The seven replicons of the *Rhizobium* genome are displayed on the outer circle of this circos visualization(*24*). Genes that were beneficial in a single niche (green inner band) or across multiple niches (generalist genes; blue inner band) and are marked for each replicon. Jitter was added along the y-axis (height) position of the circles to aid visualization of genes in close proximity.

## Plasmids are depleted in beneficial genes

To understand the benefits of plasmid and chromosomal genes, we calculated the fraction of plasmid and chromosomal genes that were beneficial in each niche. Crucially, the proportion of plasmid genes with beneficial effects on fitness was low relative to the chromosome in all niches, challenging the ecological importance of plasmids (Figure 2A).

**Figure 2:**
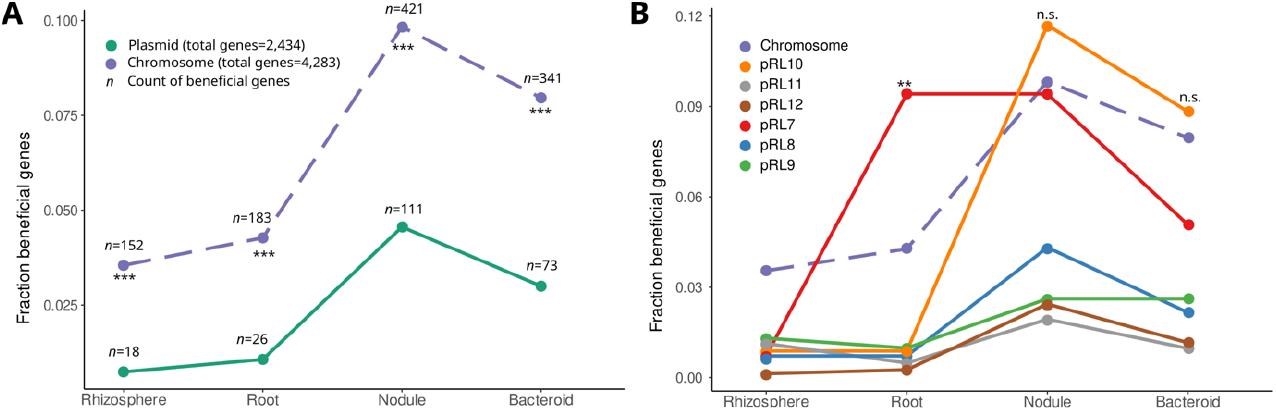
Plasmid are depleted in beneficial genes. Plots show comparisons of the prevalence of beneficial genes (ie beneficial genes/total genes) between the chromosome and all plasmid genes (**A**) and between individual replicons (**B**). We compared the proportions of beneficial genes on plasmids and the chromosomes using a normal approximation to the binomial distribution. All comparisons between plasmids and the chromosome in **A** were statistically significant under a two-tailed null hypothesis with P<1×10^−10^. In **B** we tested for an increased prevalence of beneficial genes on plasmids compared to the chromosome. Plasmid pRL7 was enriched in root adaptive genes compared to the chromosome (P one -tailed=.0017, **).

However, a limitation of this analysis is that it treats plasmid genes as a collective. If plasmids are key drivers of niche adaptation, then individual plasmids might be associated with genes involved in specialization on distinct niches. Consistent with this idea, we found two clear examples of niche-associated plasmids. Plasmid pRL10 carries genes that play important roles in the establishment of symbiotic interactions with legumes, including nitrogen fixation (*17, 21*). As expected, this plasmid was associated with genes that were beneficial during nodulation and bacteroid formation. Second, plasmid pRL7 was associated with genes that were beneficial across all of the niches associated with plants, including root colonization. Although these examples highlight the association between plasmids and niches, it is important to emphasize that plasmids were not enriched in niche-adaptive genes compared to the chromosome, except for a single case of genes involved in root specialization on plasmid pRL7.

## Plasmids are associated with niche specialist genes

If evolutionary processes drive beneficial genes to become localized to the chromosome, genes that are under strong selection should be more likely to be associated with the chromosome compared to genes that are under weak selection(*18*). To test this prediction, we compared the distribution of genes that were beneficial in a single niche (specialist genes) with those that were beneficial across multiple niches (generalist genes). The underlying assumption of this test is that genes that are beneficial in a single niche are under weak selection compared to genes that are beneficial across multiple niches when selection is considered across the entire life cycle of *Rhizobium*.

Overall, plasmids were not enriched in specialist genes compared to the chromosome (Figure 3A). However, plasmids pRL10 and pRL7 were enriched in specialist genes, reflecting the roles that these plasmids play in interactions between *Rhizobium* and plants (Figure 3C). The overall lack of specialist genes on plasmids was driven by the fact that the remaining plasmids were depleted in niche specialist genes, and this depletion was most obvious for plasmids pRL11 and pRL12. In contrast, generalist genes that were beneficial across multiple niches were strongly associated with the chromosome (Figure 3B). None of the plasmid replicons were enriched in generalist genes, and the depletion of generalist genes was particularly clear for plasmids pRL9, pRL11 and pRL12 (Figure 3D).

**Figure 3:**
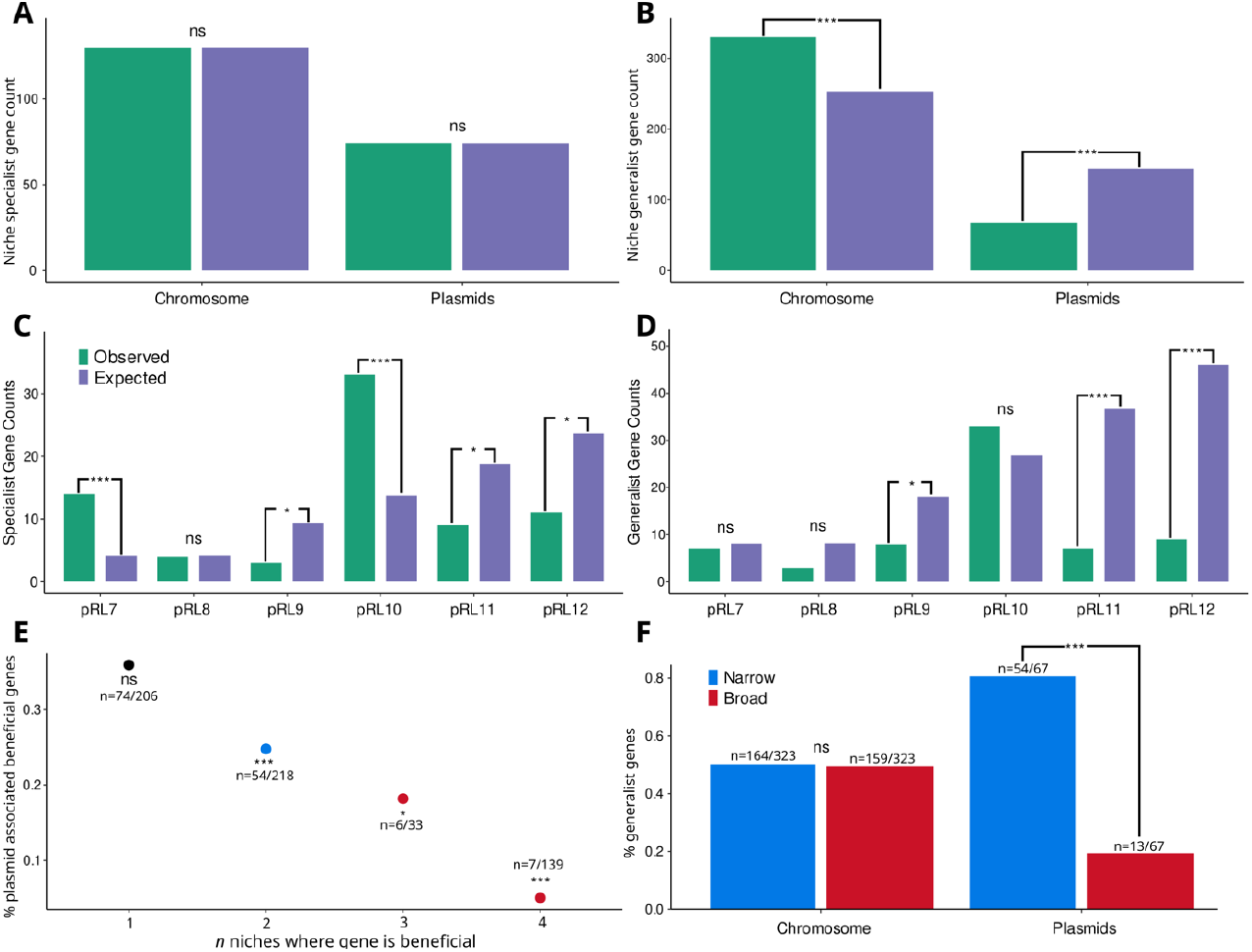
Plasmids are associated with niche specialist beneficial genes. Bar charts show the observed and expected number of specialist (A,C) and generalist (B,D) genes across the genome. Panels A and B show a comparison of all plasmid genes with the chromosome, and Panels C and D show individual plasmid replicons, with observed gene counts shown in green and expected gene counts shown in dark blue. Expected gene numbers were calculated based on the number of genes on each replicon under the null hypothesis that the prevalence of beneficial genes is equal for all replicons. We tested for beneficial gene enrichment using two-tailed binomial tests comparing all plasmids and the chromosome (Panel A,C) or individual plasmid replicons (Panel B,D). Statistical tests for individual replicons were corrected for multiple testing using the Bonferonni correction. Panel E shows the proportion of beneficial genes associated with plasmids as a function of the number of niches where the gene was beneficial. The number of plasmid associated beneficial genes are shown and we tested the null hypothesis that beneficial genes are evenly distributed across the genome using two-tailed binomial tests. Panel F shows the proportion of generalist genes that increased fitness in a narrow range (2 niches) or a broad range (3 or 4 niches) of niches for plasmids and the chromosome. We tested for a difference in the proportion of narrow and broad range generalist genes usign a normal approximati n to the binomial distribution. Significance: n.s: no significant enrichment; *, P<.05; **, P<.01; ***, P<.001;.

To further test the hypothesis that genes under strong selection become associated with chromosomes, we treated the number of niches where genes were beneficial as an ordinal variable (i.e. 1-4 niches) as opposed to a binary variable (i.e. specialist or generalist). As expected, plasmids were depleted in genes that were beneficial across multiple niches compared to the chromosome (Figure 3E). An alternative way to visualize this result is to compare the prevalence of narrow range generalist genes that were beneficial in 2 niches with broad range generalist genes that were beneficial in 3 4 niches (Figure 3F). Almost all of the generalist genes carried by plasmids were narrow range, while broad and narrow range generalist genes were equally represented on the chromosome.

## Plasmids lose beneficial genes over time

If selection favours the movement of beneficial genes from plasmids to the chromosome, then recently acquired plasmids should be rich in beneficial genes compared to ancient plasmids. Plasmids pRL9, pRL11 and pRL12 lack motility systems and have a nucleotide composition that matches the chromosome, suggesting that they were acquired by *Rhizobium* in the distant past(*17*). The remaining plasmids (pRL7, pRL8, pRL10) have divergent nucleotide composition from the chromosome and plasmid mobilization systems (pRL7 and pRL8), implying that they have been more recently acquired. To test this hypothesis, we compared the prevalence of all beneficial genes between recently acquired and ancient plasmids (Figure 4). We did not distinguish between specialist and generalist genes in this analysis, due to the fact that plasmids carried few generalist genes that were typically beneficial in only 2 niches (Figure 3E,F). Beneficial genes were over-represented on recently acquired plasmids, whereas beneficial genes were strongly depleted from ancient plasmids, suggesting that plasmids become gradually depleted in beneficial genes over time.

**Figure 4:**
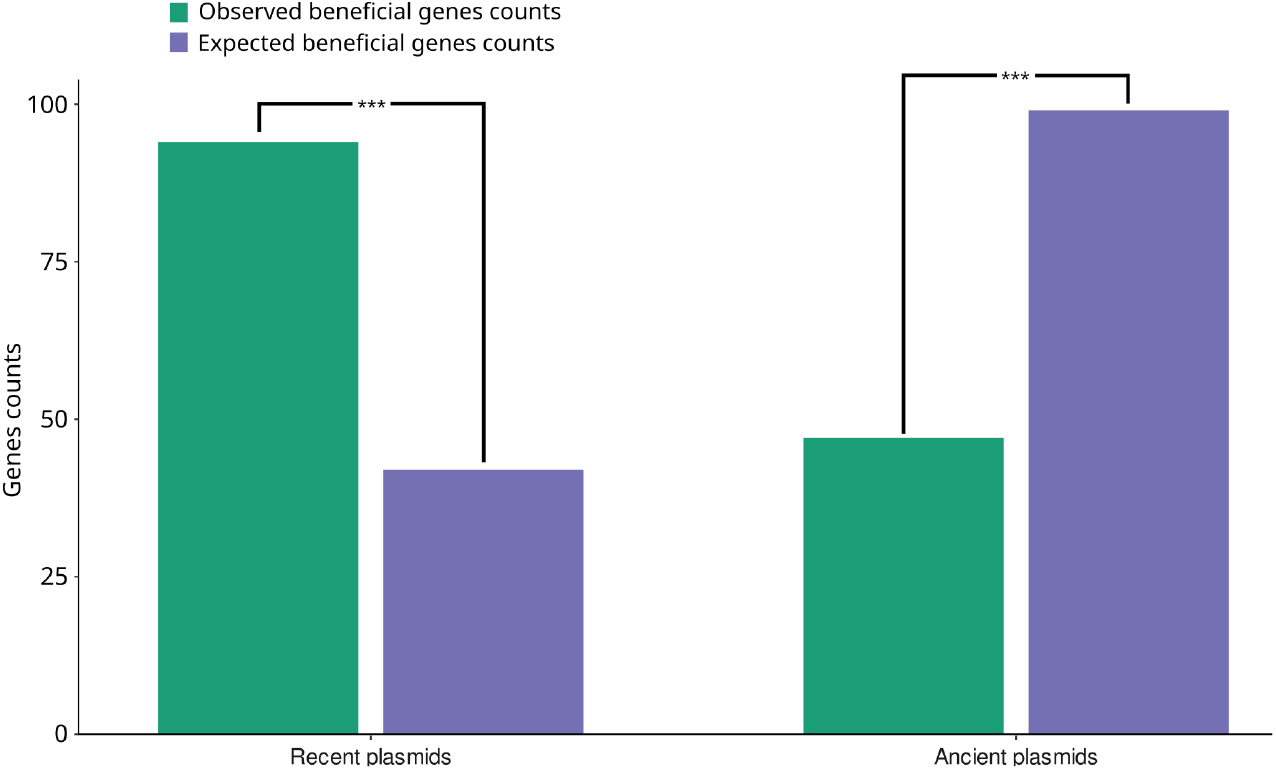
Ancient plasmids are depleted in beneficial genes. Bar charts show the expected and observed number of beneficial genes for recently acquired (pRL7,pRL8, pRL10) and ancient (pRL9,pRL11,pRL12) plasmids. Expected gene numbers were calculated based on the number of genes on each replicon under the null hypothesis that the prevalence of beneficial genes is equal across plasmids. We tested for significant deviations from expected gene counts using a two-tailed binomial test, and both P values were highly significant (P<6×10^−4^).

## Discussion

Plasmids are a ubiquitous component of bacterial genomes, but their role in adaptation remains unclear. Classic evolutionary models allowing movement of genes between chromosomes and plasmids predict that, over time, beneficial genes will become localized to the chromosome(*2*). Consistent with this model, we found that plasmids were depleted in beneficial genes compared to the chromosome (Figure 2A), because genes that were beneficial across multiple ecological niches were strongly localized to the chromosome (Figure 3). If the chromosome effectively captures beneficial genes, then we would expect plasmids to undergo a process of gradual ecological decay due to the loss of beneficial genes. Consistent with this idea, we found that ancient plasmids were depleted in beneficial genes compared to recently acquired plasmids (Figure 4). Our results suggest that plasmids acquisition provides bacteria with beneficial genes, but the movement of beneficial genes to the chromosome causes plasmids to degrade towards ecological redundancy, emphasizing the challenge of understanding how plasmids can persist over the long term (*5, 6, 25-29*).

The paradigm that plasmids play a key role in adaptation by providing bacteria with genes that are beneficial in specific ecological niches is deeply ingrained in microbiology(*9-13*). As expected from this paradigm, we found that plasmids were associated with genes that increased fitness in specific ecological niches (Figure 2B). One of the key insights from our study is that this association arises because evolution drives strongly beneficial genes, such as those that increase fitness across multiple niches, to become localized to the chromosome, leaving plasmids associated with niche specialist genes (Figure 3). We argue that this link between plasmid degeneration and niche specialization reconciles the adaptationist view of plasmids that has emerged from empirical studies with evolutionary models that predict the degeneration of plasmids.

Many of the most important forms of antibiotic resistance have been driven by the acquisition of plasmids carrying antibiotic resistance genes (*14, 30*). Our findings predict that the strong selective pressures caused by the continued large-scale use of antibiotics will stabilize resistance by accelerating the integration of resistance genes into the chromosomes of pathogenic bacteria, as has already been observed for some resistance genes(*31-33*).

## Methods

The supplementary data (Dataset S01 and Table S1-S8) was downloaded from Wheatley *et al* (*21*) where a large-scale transposon insertion sequencing experiment was conducted to identify genes required in *R. leguminosarum* bv. *viciae* 3841 (Rlv3841) to engage in symbiosis with the legume host pea (*Pisum sadvum*). This dataset listed the *R. leguminosarum* genes predicted to be required for fitness across four stages of symbiosis: (1) growth in the rhizosphere, (2) root colonisation, (3) nodulation, and (4) bacteroid formation. In Wheatley *et al* (*21*), a Hidden Markov Model was applied to classify genes into one of four state classifications based on their read-mapping statistics: essential (ES; no or very few insertions, i.e. insertions are not tolerated), defective (DE; significantly fewer insertion read counts along a significant consecutive stretch of insertion sites, i.e. insertion mutations impair growth), advantaged (AD; significantly higher insertion read counts along a significant consecutive stretch of insertion sites, i.e. insertion mutation enhances fitness), and neutral (NE; within the boundaries of a mean parameter of insertion read counts, insertion mutation has a neutral impact on fitness). To test the validity of their tn-seq experiment, Wheatley et al tested the roles of 15 genes in follow-up experiments using independently constructed mutants. Only a single one of these mutants did not recapitulate its predicted phenotype inferred from the sequencing of pooled populations of transposon mutants.

For our analysis, we used the gene lists defined in Wheatley *et al* (*21*) as being required for engaging in symbiosis (Figure 1B) which are composed of genes which were all identified with a NE classification in the input library and either an ES or DE classification in at least one of the symbiosis output libraries (rhizosphere growth, root colonisation, nodulation or bacteroid formation). As such, we are analysing genes that can be considered beneficial genes for plant-associated growth and symbiosis, as their mutation has a negative impact on fitness. We used these previously defined lists (*21*) with the additional downstream removal of genes with potential gene duplications in the Rlv3841 genome from the analysis(*17, 34*). This was used to calculate a total number of genes on each replicon as the denominator for the enrichment analysis, by subtracting potential gene duplications from the input library from the total gene numbers on the replicons. This had minimal impact on the output results.

## Supporting information

Supplementary data 1 beneficial genes

Supplementary data 2 data summaries for figures

## Acknowledgements

We thank Professor Philip Poole for his comments on an early version of this data analysis, and we thank Stu West, Liam Shaw, Michael Brockhurst and Alvaro San Millan for feedback on a draft manuscript. Figures within this publication were created using Biorender.com (Figure 1A, Figure 1B).

## Author contributions: (CREDIT system)

Conceptualization: RCM, RMW

Methodology: RCM,RMW,CL

Investigation: RCM, RMW

Visualization: CL,RMW,RCM

Funding acquisition: RCM,RW,CL

Writing - original draft: RCM

Writing - review & editing: RCM, RW,CL

## Data and materials availability

This study used publicly available datasets downloaded from Wheatley *et al* (*21*) (Dataset S01 and Table S1-S8).Data sets used in this analysis are given in Supplementary data file 1, and raw data for figures in supplementary data file 2.

